# The frequency-dependent effect of electrical fields on the mobility of intracellular vesicles in astrocytes

**DOI:** 10.1101/2020.05.22.111286

**Authors:** Yihua Wang, Thomas P. Burghardt, Gregory A. Worrell, Hai-Long Wang

**Author notes:** Correspondence to: Hai-Long Wang, Ph.D.; Gregory A. Worrell, System Electrophysiology Laboratory, Neurology Department, Mayo Clinic, Rochester, Minnesota 55905.

## Abstract

Slow-wave sleep, defined by low frequency (<4 Hz) electrical brain activity, is a basic brain function affecting metabolite clearance and memory consolidation. Although the origin of low-frequency activity is related to cortical up and down states, the underlying cellular mechanism of how low-frequency activity becomes effective has remained elusive. We applied electrical stimulation to cultured glial astrocytes while monitored the trafficking of GFP-tagged intracellular vesicles using TIRFM. We found a frequency-dependent effect of electrical stimulation that electrical stimulation in low frequency elevates the mobility of astrocytic intracellular vesicles. We suggest a novel mechanism of brain modulation that electrical signals in the lower range frequencies embedded in brainwaves modulate the functionality of astrocytes for brain homeostasis and memory consolidation. This finding suggests a physiological mechanism whereby endogenous low-frequency brain oscillations enhance astrocytic function that may underlie some of the benefits of slow-wave sleep and highlights possible medical device approach for treating neurological diseases.

## Introduction

Mounting evidence suggests that sleep is a basic brain function promoting memory-consolidation and brain metabolite clearance (1–7). Sleep is commonly classified into rapid eye movement (REM) and non-rapid eye movement (NREM) stages that are characterized by the frequency and pattern of brain electrical activity, called the electroencephalogram (EEG). NREM sleep includes three stages; with stage 3 being called slow-wave sleep (SWS) because it is characterized by slow-wave activity (SWA). SWA occurs when cortical neurons oscillate at low-frequency (1 – 4 Hz) between a hyperpolarized down-state and neuronal silence and a depolarized up-state and active neuronal firing (8).

Several independent lines of evidence support the notion that the effect of sleep on metabolite clearance is associated with brain waves at certain frequencies ranges. First, the rate of β-amyloid (Aβ) clearance in mice is improved during sleep and this improvement is associated with an increase in the prevalence of slow waves in sleep (5); Second, the Aβ/tau levels in the cerebrospinal fluid (CSF) are inversely correlated with SWA or the percentage of SWS in total sleep time (9, 10); Third, the Aβ/tau levels in the interstitial fluid (ISF) significantly increase during sleep deprivation (SD) versus sleep and the more SWA disruption causes the greater increase of CSF Aβ levels (7, 11–13); Lastly, slow waves are coupled to blood oxygen level-dependent signals and pulsatile CSF oscillations during NREM sleep (4), and the pulsatile CSF oscillations increase brain fluid mixing and diffusion in the fluid. Thus the SWS brain state defined by SWA shows improved clearance of brain waste products (14–18) and is also important in memory consolidation (19, 20). On the other hand, low-frequency EEG oscillations in SWS reflect the underlying transitions between cortical up and down states (21). However, whether the endogenous slow-waves recorded with EEG have any direct mechanistic role in metabolite clearance and memory consolidation has remained elusive.

The glial astrocyte has been selected as a targeting subject to study the effect of brain waves on metabolic homeostasis and memory consolidation because, in the central nervous system (CNS) astrocytes are believed to play an important role removing excess of toxic waste, recycling neurotransmitters, and other molecules, transporting major ions, modulating neuronal excitability, and promoting synaptic remodeling (22–28). Cellular functions of astrocytes can be monitored by their activities in intracellular vesicles. Various types of intracellular vesicles serve a variety of different functions (29), such as transporting proteins and other materials within a cell, carrying compounds (enzymes, hormones, and carbohydrates) to be secreted, and involving digestion and waste removal. Potokar and others previously reported that intracellular vesicles in astrocytes exhibit two types of mobility, directional or non-directional (30). The directional movement appears almost rectilinear, whereas the non-directional movement moves in Brownian motion.

Here, we utilized total internal reflection fluorescence microscopy (TIRFM) to study the frequency-dependent effect on the mobility of astrocytic intracellular vesicles. We applied electrical stimulation within a range of frequency (2, 20, 200Hz) on cultured rat astrocytes expressing GFP-labelled membrane protein (VAMP3 or CD63), and found that the mobility of intracellular vesicles in both directional and non-directional movement increased more than 20% under 2 Hz electrical stimulation, but remained unchanged under higher frequencies (20 Hz or 200 Hz) of electrical stimulation. This finding indicates that only slow waves elevate the intracellular activity of astrocytes, suggesting a novel glial mechanism of electrical brain activity that the functionality of astrocytes in brain homeostasis and memory formation can be directly modulated by electrical signals embedded in endogenous low-frequency brainwaves during SWS.

## Results

### Low-frequency electrical stimulation enhances the mobility of intracellular vesicles in glial astrocytes

To address the frequency-dependent effect of electrical stimulation on the mobility of intracellular vesicles, we selected three different frequencies (2 Hz as the low, 20 Hz as the intermediate and 200 Hz as the high) of electrical stimulation, and compared resulting movements of vesicles to that obtained when no electrical stimulation was applied. Each experiment was taken following the same timeline: two minutes of pre-stimulation, five minutes stimulation, and then another five minutes of post-stimulation. TIRF samplings were all collected in a one-minute duration. Two consecutive samplings were performed before the stimulation; during the 5-minute stimulation we collected three samplings separated by two one-minute gaps to avoid over bleaching; and then we recorded four consecutive samplings one minute later when electrical stimulation was complete. We observed directional and non-directional movements of intracellular vesicles in astrocytes, similar to the previous report from Potokar et al. (30). The mobility of non-directional vesicles can be described well as free Brownian diffusion and the tracks of directional vesicles appear almost rectilinear (Figure 1).

**Figure 1.**
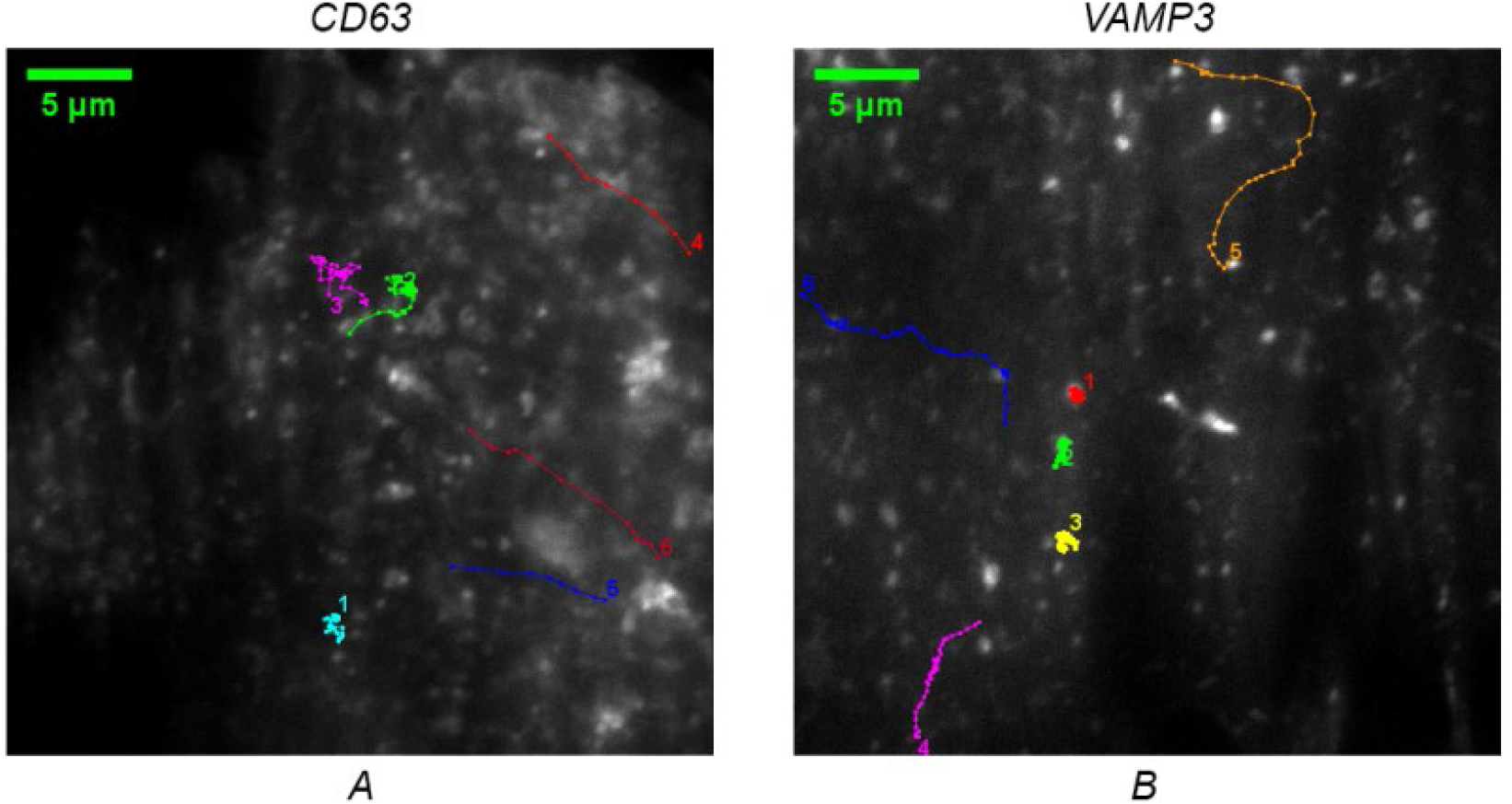
Trajectories of EGFP-CD63 (A) and EGFP-VAMP3 (B) labeled vesicles. Track 1, 2, and 3 are the trajectories of vesicles in non-directional movement. Track 4, 5, and 6 are the trajectories of vesicles in directional movement.

Results are presented in a normalized rate that compares every velocity measurements to a common scale achieved by averaging two starting samplings from each experiment. For VAMP3 labeled vesicles, the average velocities of two starting samplings were 1.03 ± 0.06 μm/s (non-directional) and 1.27 ± 0.06 μm/s (directional). For CD63 labeled vesicles the average velocity of two starting samplings were 1.24 ± 0.06 μm/s (non-directional) and 1.29 ± 0.05 μm/s (directional), respectively. Normalized rates of average velocities are illustrated in Figure 2 (non-directional) and Figure 3 (directional).

**Figure 2.**
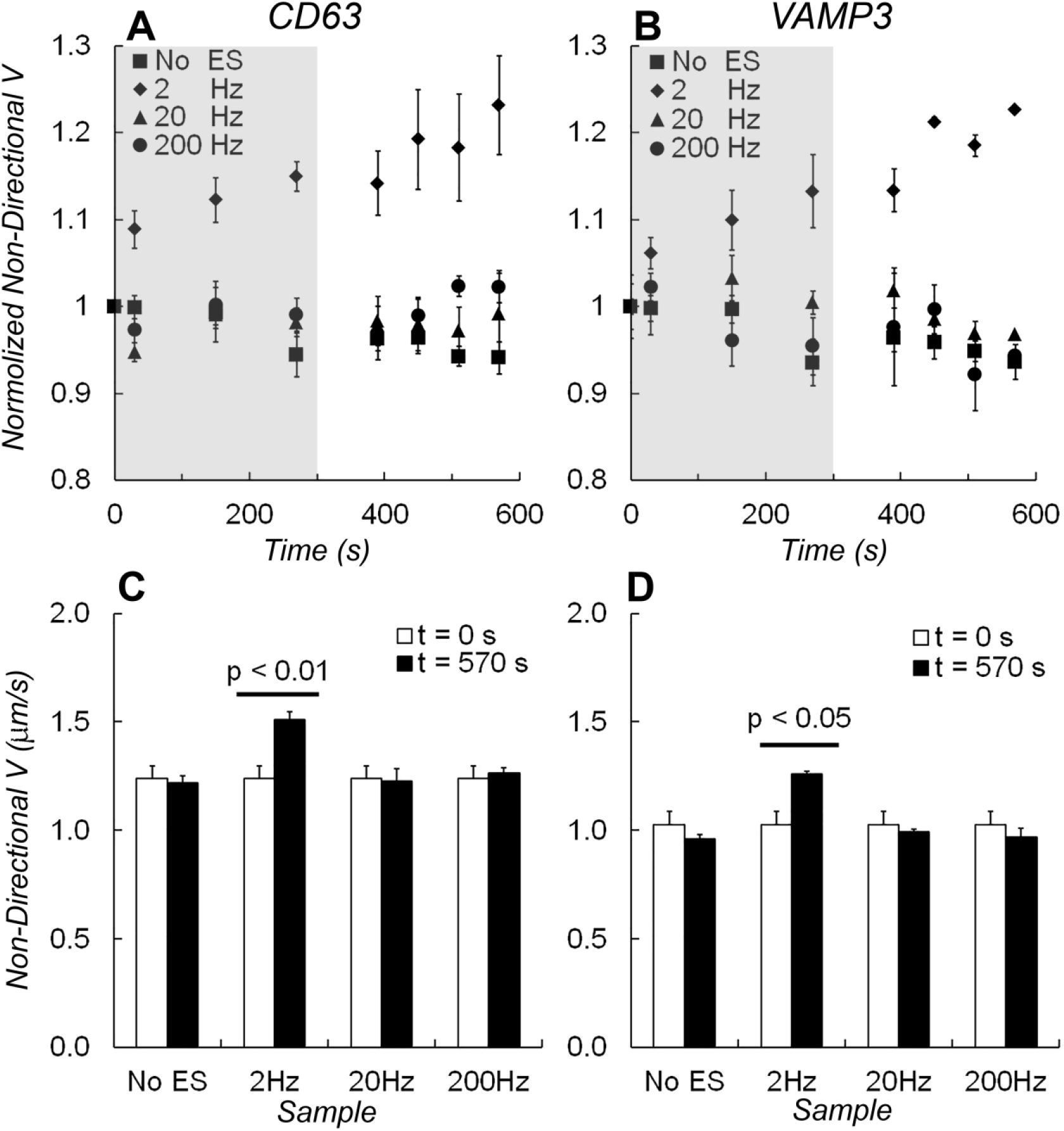
Average velocities of non-directional movement V_non-d_. The normalized velocities for non-directional movement of **A)** EGFP-CD63 and **B)** EGFP-VAMP3 labeled vesicles during the experiment timeline. Electrical stimulations were applied at 2 Hz (diamonds), 20 Hz (triangles), 200 Hz frequency (circles), or without stimulation (squares). The grey shades indicated the time when electrical stimulation was applied. **C)** A bar graph shows changes of V_non-d_ measured before (at t = 0 s) and after (at t = 570 s) electrical stimulations from EGFP-CD63 labeled vesicles. **D)** A bar graph shows changes in V_non-d_ measured from EGFP-VAMP3 labeled vesicles. One-way ANOVA test showed that the average velocity of non-directional vesicles changed significantly after 2 Hz electrical stimulation (p < 0.01 in EGFP-CD63 data and p < 0.05 in EGFP-VAMP3 data). The changes in velocity caused by electrical stimulation at 20 Hz and 200 Hz frequency, or without stimulations were insignificant.

**Figure 3.**
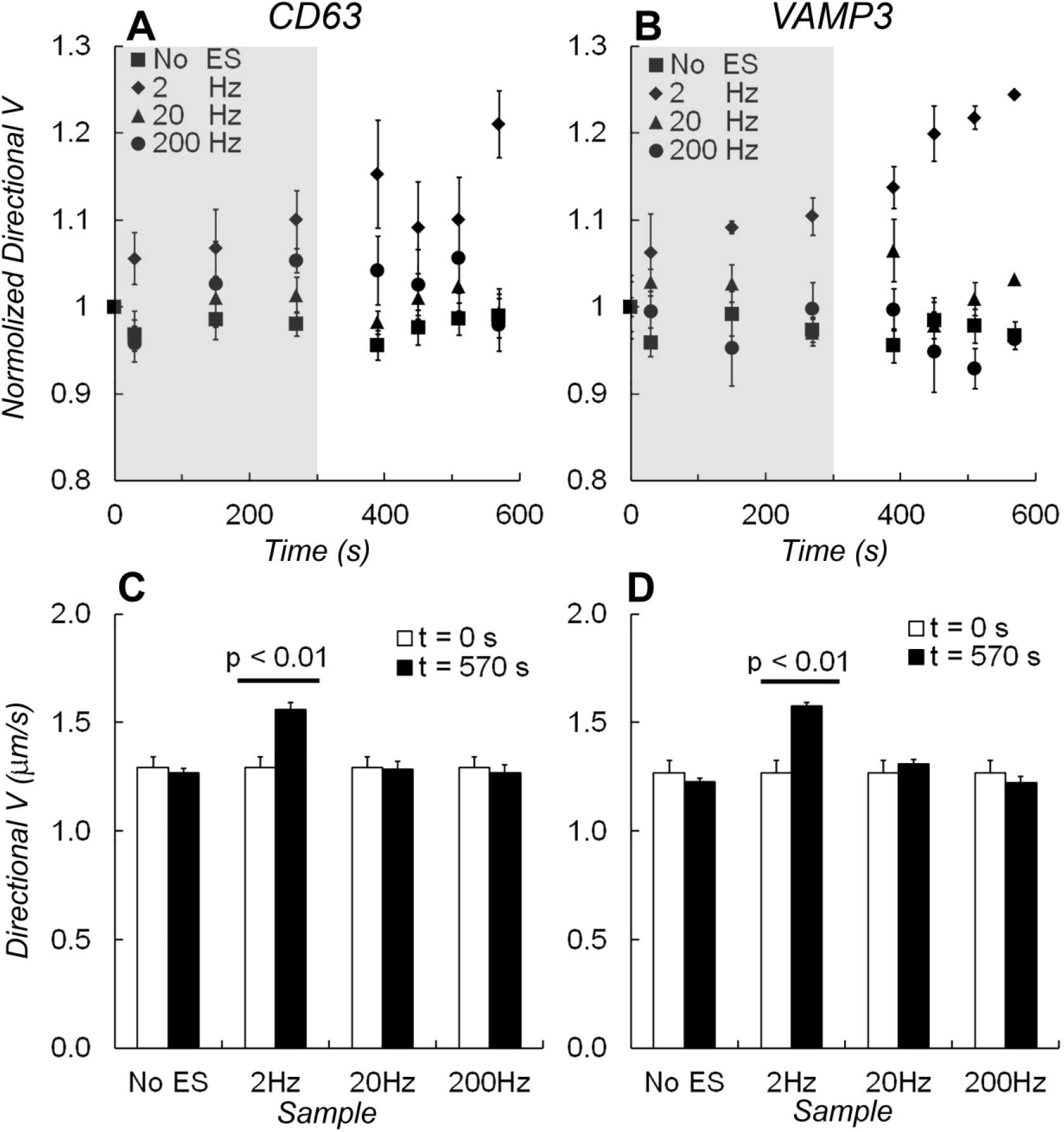
Average velocities of directional movement V_d_. The normalized velocities for directional movement of **A)** EGFP-CD63 and **B)** EGFP-VAMP3 labeled vesicles during the experiment timeline. Electrical stimulations were applied at 2 Hz (diamonds), 20 Hz (triangles), 200 Hz frequency (circles) or without stimulation (squares). The grey shades indicated the time when electrical stimulation was applied. **C)** A bar graph shows changes of V_d_ measured before (at t = 0 s) and after (at t = 570 s) electrical stimulations from EGFP-CD63 labeled vesicles. **D)** A bar graph shows changes of V_d_ measured from EGFP-VAMP3 labeled vesicles. One-way ANOVA test showed that the average velocity of non-directional vesicles changed significantly after 2 Hz electrical stimulation (p < 0.01). The changes in velocity caused by electrical stimulation at 20 Hz and 200 Hz frequency, or without stimulations were insignificant.

For non-directional vesicles, the average velocity (V_non-d_) increased immediately after 2 Hz electrical stimulation started and kept increasing linearly during the course when stimulation was applied. This increase of velocity stopped 2 ~ 3 minutes later after the stimulation ended and stayed unchanged till the end of data collection. With 2 Hz electrical stimulation, the total increase of V_non-d_ was 23% for CD63 labeling vesicles (Fig 2A) and 21% for VAMP3 labeling vesicles (Fig 2B), respectively. One-way ANOVA test comparing measurements made before and after electrical stimulation showed that the increase was significant (p < 0.01 in CD63 data and p < 0.05 in VAMP3 data). However, without electrical stimulation or with 20 Hz or 200 Hz electrical stimulations, changes in velocities for both CD63- and VAMP3-labeling vesicles are insignificant (Fig. 2C, 2D). Two-way ANOVA test showed that V_non-d_ in 2 Hz electrical stimulation measurement was increased compared to others (p < 0.01) and no significant difference was found when compared between those with no stimulation to 20 Hz or 200 Hz electrical stimulations.

For directional vesicles, the average velocity (V_d_) was increased by 20% for CD63 labeled vesicles and 22% for VAMP3 labeled vesicles (Fig. 3A, 3B) under 2 Hz electrical stimulation (p< 0.01). There was no significant change in V_d_ with no stimulation and with 20 Hz or 200 Hz electrical stimulation (Fig. 3C, 3D).

## Discussions

In this work, we investigated the frequency-dependent effect of externally applied electrical stimulation on the trafficking of astrocytic intracellular vesicles. Using EGFP tagged membrane proteins (CD63 and VAMP3), we found that the low frequency 2 Hz electrical stimulation increases the mobility for both directional and non-directional vesicles by more than 20%, but both the intermediate (20 Hz) and high (200 Hz) frequency electrical stimulation has no effect, just as when no stimulation was applied. We concluded that the intracellular activity of astrocytes can be elevated by electrical slow waves.

CD63 and VAMP3 reside in various types of intracellular vesicles and are involved in different processes of substance degradation and transportation. CD63 belongs to the tetraspanins superfamily of surface-associated membrane proteins, and mainly resides in lysosomes and multivesicular bodies (MVBs), with a small pool on the cell surface (31). After being transported to the plasma membrane, the majority of MVBs fuse with the cell membrane and release their contents called exosomes to the extracellular space. Some MVBs fuse with lysosomes for degradation (32). CD63 is one of the three biomarkers that predominantly localize on exosomes. CD63 containing vesicles in astrocytes are involved in the regulation of ATP release (33). VAMP3 is an essential component of the SNARE complexes (soluble N-ethylmaleimide-sensitive factor attachment protein receptor) that participate in the fusion of various vesicles to the cytoplasmic face of membranes. VAMP3 can be found in early endosomes, recycling endosomes, and intermediate transport vesicles (34–38). VAMP3 containing vesicles in astrocytes participate in the recycling of plasma membrane glutamate transporters and in modulating the efficacy of glutamate uptake (39).

The increased mobility of intracellular vesicles caused by 2 Hz electrical stimulation is likely nonspecific to CD63 or VAMP3. Low-frequency oscillations in electrical fields may improve the mobility of vesicles carrying other molecules as well, including neurotransmitters (adenosine triphosphate (ATP)), neurotransmitter transporters (glutamate transporters), neuromodulators (D-serine), growth factors (brain-derived neurotrophic factor), hormones and peptides (atrial natriuretic peptide, (ANP)), and the specific water channel of aquaporin-4 (33, 39–44). These substances secreted by astrocytes contribute to CNS homeostasis and development.

Astrocytes release their substances through several distinct pathways, including diffusion through plasma membrane channel, translocation by multiple transporters, and regulated exocytosis (29). The mobility of intracellular vesicles is a limiting fact controlling the substance release (45). Thus, increased mobility of intracellular vesicles leads to an enhanced release of astrocytic cargo and therefore affects CNS homeostasis and development.

Previous studies have shown that intracellular vesicles in astrocytes exhibit two kinds of movement: the directional movement relies on cytoskeleton and motor proteins, whereas the non-directional movement is driven by diffusion (30, 46–50). The rate of vesicle diffusion is determined by hydrodynamic interactions, vesicle collisions, cytoplasm viscosity, and other factors (51). It is reasonable to assume that the diffusional transport of solutes within cells could also be enhanced by electrical slow waves, which effectively enhances the rates of astrocytes intracellular biochemical processes (52).

Our finding that the low-frequency electrical stimulation, not the intermediate or high frequency, elevates the intracellular activity in astrocytes is a possible mechanism by which the endogenous SWA of SWS supports cellular homeostasis. The brainwave frequencies vary in a wide range between sleep-wake cycles. The effect of sleep on metabolite clearance is stronger during the most restorative deep sleep stage with more low-frequency (< 4 Hz) EEG oscillations (5–7, 9–13).

Astrocytes are important to remove the excess of toxic waste in the CNS, maintain neurotransmitters homeostasis, regulate the extracellular space volume, transport major ions, and regulate synaptic connectivity and transmission (22–24). Particularly in the glymphatic pathway (14, 15) that enables exchange between the para-arterial space and the interstitial space and allows waste products to be removed away from the arteries to the veins. This kind of exchange is facilitated by the AQP4 water channels presented on the astrocytic endfeet facing the perivascular space. Fluid and solutes pass through perivascular space either via the AQP4 water channels or the intercellular clefts (14, 15, 53). A recent functional magnetic resonance imaging study demonstrated that pulsatile CSF oscillations inflow to brain ventricles every 20 s during SWS (4), which is possibly caused by the pulsations of blood volume in the brain and facilitates the clearance of waste products during SWS. Also, the clearance rate of Aβ and the interstitial space volume in mouse brain increase during sleep associated with an increase in the prevalence of slow waves (5). Our finding on the elevation of intracellular activity in astrocytes induced by low-frequency electrical stimulation may explain why this increase of waste clearance occurs in SWS.

SWS is also important in memory consolidation, particularly for long-term memory (19, 20). In older adults, a decline in declarative memory consolidation is found to be associated with the decrease of SWA in NREM (54–61) but can be causally reversed by experimental enhancement of SWA in NREM (62–64). It is worth noting that astrocytic glycogen metabolism and learning-induced astrocytic lactate release are required for long-term memory formation (27). It is the activation of astrocytes, but not the increase of neuronal activity alone (28), which results in enhanced memory allocation and cognitive performance. Our results provide a possible cellular mechanism for how astrocytes could mediate long-term memory formation (25–28, 65) during SWS.

The molecular mechanism for the frequency-dependent effect of electrical stimulation on the mobility of intracellular vesicles remains unclear. Cytoskeleton and motor proteins could be involved in the directional mobility, whereas the diffusion in local microenvironments of cytoplasm might affect the non-directional movement of vesicles (30, 47–50, 66). Different protein carrying vesicles could behave differently. For example, a previous study shown that excessive ATP can trigger the elevation of intracellular Ca^2+^ concentration in astrocytes due to increased mobility of vesicles containing vesicular glutamate transporter 1 but also to reduced mobility of vesicles containing atrial naturetic peptide (67). It will be interesting to test how the frequency of electrical stimulation affects motor proteins and cytoskeleton. The traffic deficiency of astrocytic intracellular vesicles is associated with many neurological diseases, including Alzheimer’s disease, Rett syndrome, and amyotrophic lateral sclerosis (68–71). Using electrical stimulations to enhance the traffic of intracellular vesicles may prove to have therapeutical benefits.

Besides the mobility of intracellular vesicles, gene expression in astrocytes can also be affected by the sleep-wake cycles (72). Previously, we reported a frequency-dependent effect of externally applied electrical stimulation on the secretion of astrocytic extracellular vesicles (EVs), including the profiles of EV size distribution, EV surface proteins, and EV-carrying microRNAs (73).

Unlike other studies of electrical stimulation on cultured astrocytes that mostly focused on the applied current amplitude (4 – 1500 mV/mm) rather than the frequency (74–78), this report, to the best of our knowledge, demonstrates for the first time that the mobility of intracellular vesicles in astrocytes can be increased only at a low-frequency electrical stimulation. This frequency-dependent effect of electrical fields on cellular functionalities likely applies to other cell types, such as neurons, microglial, or even cancer cells.

In summary, we reported here our study on the trafficking of intracellular vesicles in astrocytes under externally applied electrical stimulation at different frequencies. We found that the mobility of intracellular vesicles for both directional and non-directional movements was elevated by more than 20% with low frequency (2 Hz) electrical stimulation, but remained unchanged with electrical stimulations at an intermediate (20 Hz) or higher frequency (200 Hz). This finding of the frequency-dependent effect on astrocytes provides a unique glial mechanism that may be important for both brain homeostasis and memory consolidation, and also have therapeutic implications of medical devices to provide electrical brain stimulation. It certainly extends our understanding of the basic biology about why deep sleep is the most restorative stage of sleep. Further investigations on the molecular mechanisms of how electrical fields modulate cellular functions are needed.

## Materials and methods

### Cell cultures

Animal experiments complied with the National Institutes of Health Guide for the Care and Use of Laboratory Animals and were approved by the Mayo Clinic Institutional Animal Care and Use Committee. Pregnant Lewis rats were purchased from Harlan Sprague Dawley (Indianapolis, IN, USA). Astrocytes cultures were established from the cerebral cortices of fetal rats (embryonic days 18-21) as previously described (79). Astrocytes cultures were maintained in a growth medium composed of Dulbecco’s modified Eagle’s medium (DMEM) supplemented with 10% fetal calf serum (FCS), 1 mM sodium pyruvate, 100 IU/mL of penicillin, 100 μg/mL of streptomycin, and 292 μg/mL of L-glutamine at 37°C in a 5% CO_2_-95% air incubator. Microglia depletion was achieved by adding 100 μg/mL of Clodrosome (Encapsula NanoSciences, Brentwood, TN, USA) to the growth medium for 12 h, then replacing with fresh medium for another 48 h, and was finally confirmed by Iba1 immunostaining (FUJIFILM Wako Pure Chemical Corporation, Osaka, Japan).

### Cell transfection

Transfection was achieved using astrocytic cultures and a transfection reagent, TransIT-293 (Mirus, Madison, WI, USA). Plasmid encoding EGFP-VAMP3 was a gift from Dr. Thierry Galli (Addgene plasmid # 42308 and # 42310) (80, 81) and CD63-pEGFP C2 was a gift from Paul Luzio (Addgene plasmid # 62964). Plasmid DNA was prepared using HiSpeed Plasmid Maxi Kit (QIAGEN, Germantown, MD, USA). DNA purity and concentration were identified using NanoDrop (ThermoFisher SCIENTIFIC, Grand Island, NY, USA). The purified DNA was incubated for 20 min at room temperature with the TransIT-293 reagent in Gibco™ Opti-MEM reduced-serum medium. The mixture (130 μL) was dispersed into a 35 mm poly-D-lysine coated glassbottom dishes (P35GC-1.0-14C, MatTek, Ashland, MA, USA) of astrocytes in complete growth medium (2.5 mL). Each dish contained 1.3 μg of either EGFP-CD63 or EGFP-VAMP3. And then the astrocytes in the medium were incubated for 2-3 days at 37°C in a 5% CO_2_-95% air incubator. Cell density was 0.6 - 1.0 × 10^5^ count/dish.

### Electrical stimulation of astrocytes

Electrical stimulation was applied to culture astrocytes using a stimulus isolator (A365, World Precision Instruments, Sarasota, FL, USA) and a digitizer (Axon Digidata 1440A, Molecular Devices, San Jose, CA, USA). The current output from stimulus isolator A365 was converted to constant voltage output with a 1KΩ bridging resistor, and then connected to a perfusion insert designed for electric field stimulation on 35 mm culture dishes (RC-37FS, Warner Instrument, Hamden, CT, USA) that provides 5 mV/mm field strength in the open region of the insert. Five minutes of electrical stimulation was applied for each experiment. We selected three frequencies, 2 Hz, 20 Hz, and 200 Hz, all have 0.1 ms pulse width. To guarantee that the total number of pulses are the same during 5 minutes of stimulation, multiple gaps were inserted at 20 and 200 Hz frequencies to achieve 600 total pulses for each experiment. TIRF samplings were all collected in a one-minute duration. Two consecutive samplings were performed before the electrical stimulation, three samplings separate by two one-minute gaps during the stimulation to avoid over bleaching, and then four consecutive samplings were obtained after one minute when the electrical stimulation was complete.

### Imaging acquisition and processing

Cells were observed by using an inverted microscope (Axio Observer Z1, ZEISS, Oberkochen, Germany) equipped with total internal reflection fluorescence (TIRF) illumination (see Figure S1 in supplement) that is confined to the glass surface with an evanescent field in ~100 nm depth penetrating cells setting above the glass surface (82). An excitation wavelength of 488 nm was provided by a 100 mW argon laser. Images were captured through a 100×/1.46 alpha Plan-Apochromat oil-immersion objective (ZEISS) and a digital CMOS camera with a pixel size of 6.5 μm × 6.5 μm (ORCA-flash4.0, HAMAMATSU, Hamamatsu City, Japan) driven by ZEN imaging software. TIRF imaging frames were streamed directly to a hard drive for 60 seconds at 100 ms/frame covering a 512 ×512-pixel region of the camera chip. The complete growth medium was replaced with phosphate-buffered saline supplemented with 0.9 mM CaCl_2_ and 0.5 mM MgCl_2_ before imaging acquisition. All experiments were performed at 37°C. DiaTrack software (83) was used to localize and track movements of all detected vesicles. Based on a directionality index (a ratio comparing the maximal displacement to the total track length (30)), we categorized movements of vesicles into two groups: directional (index >= 0.5) and non-directional (index < 0.5). Custom Matlab codes were used to categorize and calculate the velocity of vesicle movements.

### Statistics

Experiments with/without electrical stimulation were repeated 4-5 times for both EGFP-CD63 and EGFP-VAMP3 transfected astrocytes. Cells from one culture dish were used only for one experiment. All data were presented as mean ± standard deviation unless specified otherwise. Significance was tested using one-way or two-way ANOVA (Tukey Kramer method). All statistical operations were performed using Matlab (MathWorks, Natick, MA, USA).

## Acknowledgments

The work was supported by an NIH BRAIN initiative R01 grant to HLW and GAW (NS112144 from NINDS and NIMH) and the Minnesota Partnership for Biotechnology and Medical Genomics (MNP #17.16). We thank Dr. Vanda A Lennon’s group who prepared rat astrocyte cultures.

## Author Contributions

H. W. designed experiments; H.W. and Y.W. carried out the experiments and analysis; Y.W., H.W., G.W., and T. B. wrote the paper.

## Supplementary Materials

**Figure S1.**
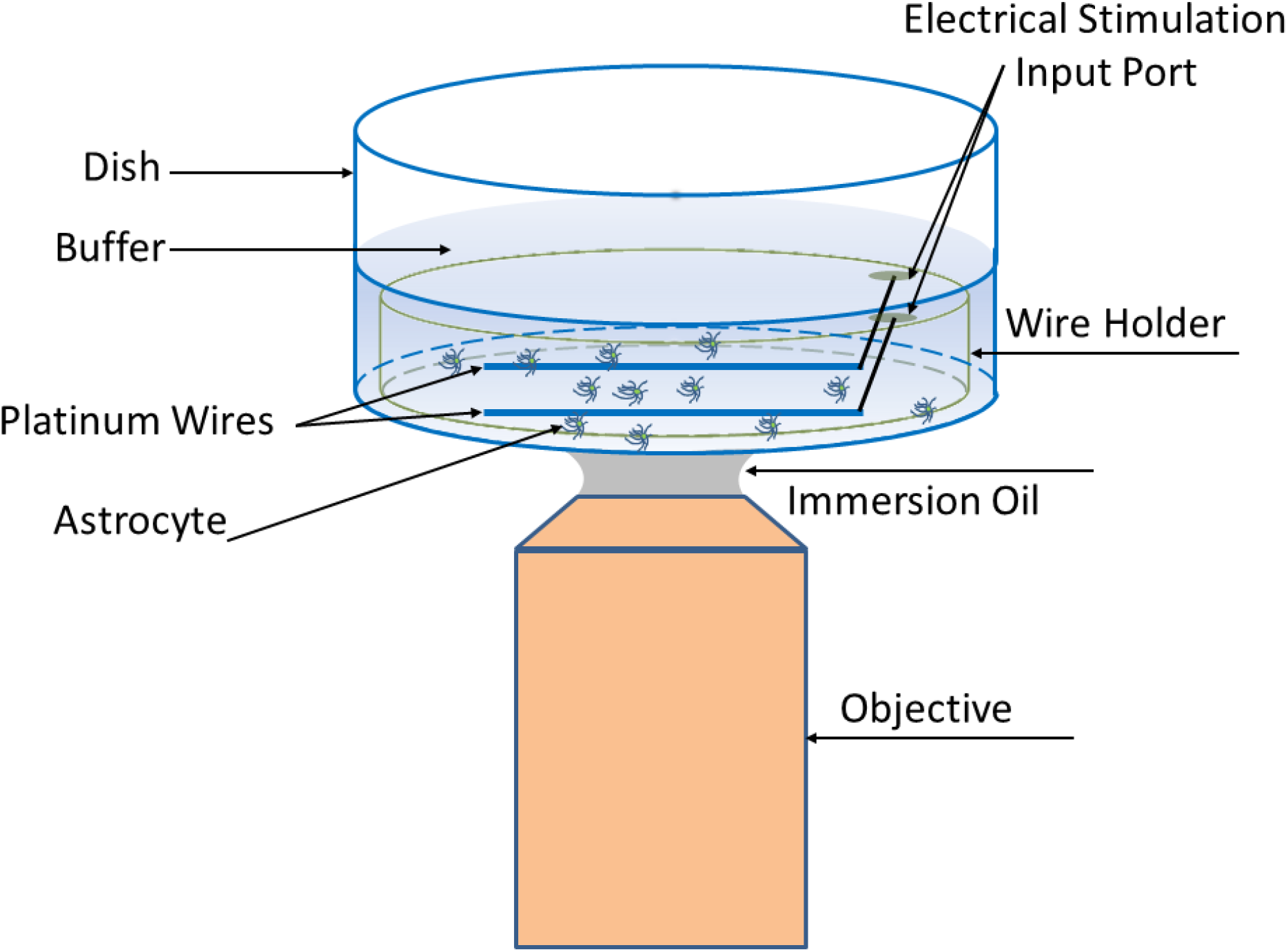
A schematic diagram of the experimental setup for measuring mobility of vesicles in astrocytes using TIRF microscope. The output of stimulus isolator was connected to platinum wires through two electrical stimulation input ports.

Movie 1 and 2 (see attachment)

